# Genomic Characterization of Novel Endophyte Strains from Tall Fescue Shows Genome Fragmentation Post-Hybridization

**DOI:** 10.1101/2025.02.07.637142

**Authors:** Kendall Lee, Philip Bentz, Justin Vaughn, Ali Missaoui

**Affiliations:** Institute of Plant Breeding, Genetics and Genomics, University of Georgia, Athens, GA 30602; Department of Plant Biology, University of Georgia, Athens, GA 30602; Genomics and Bioinformatics Research, USDA-ARS, Athens, GA 30605; Department of Crop and Soil Sciences, University of Georgia, Athens, GA 30602

## Abstract

Interspecific hybridization in fungi has gained attention for its role in fungal evolution and potential commercial applications. Successful hybridization can enhance fitness and facilitate adaptation to new ecological niches. However, the genomic consequences of hybridization in fungi remain poorly understood. *Epichloë* is a genus of fungi that includes both non-hybrid and hybrid species, with the hybrids forming through parasexual hybridization and reproducing asexually. Some *Epichloë* hybrids are of commercial significance, as they colonize *Lolium arundinaceum* (Schreb.) Darbysh., a crucial forage and turf grass species. In this study, we sought to generate high-quality genome assemblies for two previously uncharacterized *Epichloë* hybrid strains, both of which are similar to *Epichloë* sp. FaTG-3. We aimed to characterize their genomes and examine the effects of parasexual interspecific hybridization on fungal genome structure. Our results reveal that the genomes of both strains are rich in AT-rich blocks and repetitive elements. Upon comparison with putative progenitor genomes, we observed significant fragmentation and rearrangement. Despite the genomic instability, more than 85% of gene homologs from each progenitor species were retained. This study demonstrates that while parasexual hybridization dramatically alters genome structure, it does not significantly affect gene content.

## INTRODUCTION

Interspecific hybridization has long been recognized as a key driver of evolution and domestication in certain plant species (Purugganan, 2019; Soltis, Visger, & Soltis, 2014). More recently, hybridization events in fungi have garnered attention for their role in evolution and their potential commercial applications. In fungi, hybridization events have been instrumental in the evolution of plant pathogens, such as the causative agents of Dutch elm disease, and have contributed to the development of yeasts used in the production of beer, bread, and other fermented products (Brasier, 2001; Gallone et al., 2019). Newly formed fungal hybrids often exhibit genetic instability, leading to significant changes in genome structure, phenotype, transcriptome, and proteome (Steensels, Gallone, & Verstrepen, 2021). While research into hybrid fungi is emerging, many aspects of their genetic dynamics remain poorly understood. Investigating the genomic impacts of interspecific hybridization in fungi can provide valuable insights into hybrid fungi evolution and may inform advancements in artificial hybridization or genetic engineering for agricultural and commercial purposes.

Fungi can hybridize either sexually or parasexually. Parasexual hybridization, which occurs exclusively in fungi and prokaryotes, is driven by hyphal fusion events that lead to the formation of heterokaryons (multinucleate cells). These hybrids typically revert to homokaryons (a single nucleus) quickly, and the resulting genomes can display significant diversity. Parasexual hybridization may result in chromosome and gene losses, with heteroploidy sometimes being retained in asexual systems (Steensels et al., 2021). In nature, parasexual hybridization followed by clonal reproduction can enhance the fitness of certain fungal species. However, various barriers, such as geographical, temporal, and ecological isolation, must be overcome for hybridization to occur. Even after hybridization, post-hybridization barriers, including genetic incompatibility, may arise. While sterility often limits the success of sexual hybrids, clonal species can bypass this barrier. If hybridization is successful, the new hybrid may either outcompete the progenitor species due to increased fitness or occupy a different ecological niche. In this study, we explore these mechanisms using an asexual, interspecific hybrid Epichloë species.

Hybrid Epichloë fungi are mutualistic symbionts of many cool-season grasses, where they act as endophytes. These grasses, prevalent in temperate regions, are ecologically significant and commercially valuable. The presence of Epichloë endophytes can help grasses resist abiotic and biotic stressors, such as pests and drought. The endophytes produce alkaloids that contribute to these protective effects (Lee, Missaoui, Mahmud, Presley, & Lonnee, 2021). In agricultural settings, certain Epichloë species are exploited to improve host grass performance. Tall fescue (*Lolium arundinaceum*), a cool-season grass used for forage and turf, is one such grass. Endophyte-infected tall fescue is highly sought after for its persistence. The species exhibits different morphotypes, including continental, Mediterranean, and rhizomatous types, each associated with specific Epichloë species (Ekanayake et al., 2017; Young et al., 2014). The Epichloë endophyte can produce four alkaloid classes: ergot alkaloids, peramine, lolines, and indole-diterpenes (Takach et al., 2012). Peramine and lolines have demonstrated anti-insect properties, making them highly desirable, while ergot alkaloids can cause fescue toxicosis in grazing livestock. To address this, non-toxic, ergot-alkaloid-free commercial varieties of tall fescue have been developed. Biosynthetic genes for these alkaloids have been identified, with peramine being synthesized by a single gene (*perA*), while other alkaloids are produced by complex gene clusters (Hettiarachchige et al., 2019).

The Epichloë species associated with tall fescue are primarily interspecific hybrids that reproduce asexually. These hybrids form through hyphal fusion between progenitor species (Tsai et al., 1994). The progenitor species are sexual and haploid, often regarded as more pathogenic than the hybrid species. Hybridization followed by clonal reproduction enables the hybrid fungi to remain mutualistic with their host while enhancing their fitness. The hybridization of different progenitor species leads to a phenomenon known as “alkaloid pyramiding,” where the hybrid inherits alkaloid biosynthesis genes from each progenitor, resulting in increased alkaloid production (Charlton et al., 2014; Schardl, Young, Faulkner, Florea, & Pan, 2012).

Endophyte species are typically characterized by their chemotype, ploidy, host grass, and mating type. Two mating types, A and B, exist in Epichloë, with interspecific hybrids often possessing multiple mating type genes inherited from their progenitors. *E. coenophiala* is a triploid hybrid with three progenitor species, while other species such as *E. sp. FaTG-2*and *E. sp. FaTG-3* are diploid with two progenitors (Takach & Young, 2014). The progenitors of *E. coenophiala* include *E. festucae*, *E. typhina*, and an unknown *Lolium*-associated endophyte. Other species, such as *E. sp. FaTG-2*, *E. sp. FaTG-3*, and *E. sp. FaTG-4*, are the result of hybridization between different combinations of progenitors. *E. coenophiala*produces all four alkaloid classes and has a mating type of AAA, while other species may lack certain alkaloids or exhibit different mating types (Young et al., 2014). Phylogenetic classification of Epichloë species is often based on nuclear housekeeping genes such as *tubB* and *tefA*, as well as the alkaloid gene *perA*, which is more informative for studying species evolution (Ekanayake et al., 2013).

Despite the availability of chromosome-resolved reference genomes for non-hybrid Epichloë species, hybrid species remain challenging to study due to their complex genomic structure. A reference genome for *E. coenophiala* exists at the contig level, assembled using a combination of sequencing technologies, including Illumina, Roche pyrosequencing, Sanger sequencing, and Oxford Nanopore (Oxford, UK) single-molecule sequencing (Florea, Jaromczyk, & Schardl, 2021). Although valuable, this genome is not fully resolved and is less suitable for applications requiring the identification of large structural variations.

In this study, we aimed to extract previously uncharacterized endophyte strains from PI accessions of tall fescue, representing both continental and Mediterranean types, and develop high-quality genome assemblies. These assemblies were then compared to investigate the genomic impact of interspecific hybridization in Epichloë species.

## MATERIALS AND METHODS

### Endophyte material

Endophytes were obtained from tall fescue PI accessions provided by the USDA-ARS Germplasm Resources Information Network (GRIN). Endophytes were cultured from the tall fescue by surface sterilizing cross-cut pieces of tiller with ethanol and bleach and placing them on sterile petri dishes with Potato Dextrose Agar (PDA) media with added 100ug/m ampicillin, 50ug/mL streptomycin and 50ug/mL chloramphenicol and leaving them for a month in the dark at room temperature. Once the endophyte had grown out of the tiller, a subculture was removed and placed on a fresh PDA plate where it was allowed to grow for a month in the dark at room temperature (Fig. 1).

**Figure 1.**
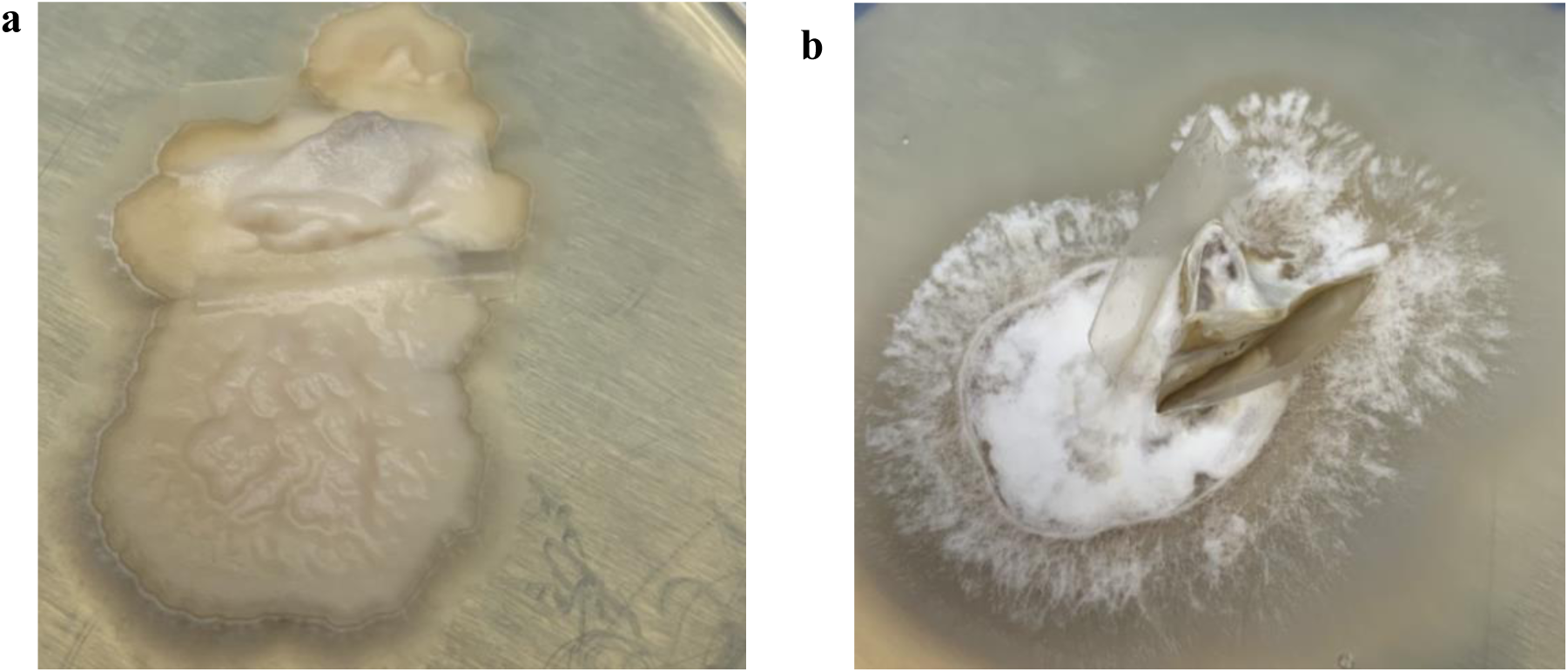
**Images of the morphology of fungal cultures of *Epichloë* strains a) 212 and b) 1033.**

### DNA Preparation and Sequencing

DNA extraction was done from fungi cultures. A piece of fungi was ground using liquid nitrogen in a sterilized mortar and pestle. The ground fungi underwent DNA extraction using a modified version of the CTAB protocol from Al-Samarrai and Schmid (2000). Initial DNA quality was checked on a NanoDrop™ 8000 Spectrophotometer. Further quality was checked using the Georgia Genomics and Bioinformatics Core’s Fragment Analyzer: High Sensitivity Large Fragment (50k). HiFi reads were generated by PacBio Sequel II SMRTCell on 1 SMRT cell for three strains (Georgia Genomics and Bioinformatics Core, Athens, GA).

### Genome Assembly and Analysis

Demultiplexed subread.bam files were converted to circular consensus sequences using the pbcromwell run pb_ccs workflow from SMRT-Link v.11.0.0 (PacBio, 2022). Prior to assembly, the raw circular consensus sequences were used to create kmer plots to analyze genome composition and sequencing quality. Jellyfish v. 2.3.0 was to create the histogram and GenomeScope 2.0 was used to plot the histograms (Marçais & Kingsford, 2011; Ranallo-Benavidez, Jaron, & Schatz, 2020) (Fig. S1). The circular consensus sequences were then assembled into contigs using Hifiasm with -l0 option to disable purging (Cheng, Concepcion, Feng, Zhang, & Li, 2021). This option was used due to the similarity of probable progenitor species. Genome assemblies were quality-checked using Quast v. 5.0.2. and BUSCO v. 5.2.2 (Gurevich, Saveliev, Vyahhi, & Tesler, 2013; Manni, Berkeley, Seppey, Simão, & Zdobnov, 2021). Assembly coverage was determined by mapping the circular consensus sequences back to the genome assembly using Minimap2 v. 2.22 and visualized in IgV v. 2.12.3 and analyzed using BEDTools v. 2.3.0 (Li, 2018; Quinlan & Hall, 2010; Robinson et al., 2011). Telomere motifs (start: taaccc, end: ttaggg) were identified from the previously assembled *Epichloë festucae* genome (ASM381444v1) and searched across all contigs in Geneious Prime (“Geneious Prime 2023 “; Winter et al., 2018). Each strain assembly had one long contig that was of low coverage, contig 1 in 212 and contig 18 in 1033. The contigs were investigated using the mito_filter.sh script from OcculterCut v. 1.1 and determined not to be mitochondrial DNA (Testa, Oliver, & Hane, 2016). Pieces of the contigs were then analyzed in BLAST and were determined to be bacterial contamination. The contigs were then removed from the assemblies.

### Species Tree

Non-hybrid *Epichloë* genomes were downloaded from NCBI. Predicted protein sequences for single-copy genes were obtained from BUSCO output for all genomes except *E. festucae*. *E. festucae* protein sequences were downloaded from NCBI. We used Orthofinder v.2.5.4 (Emms & Kelly, 2019) to infer single-copy orthologous genes shared across nine species of *Epichloë,* including strain 1033 and 212 from this study, and *Fusarium abutilonis* as an outgroup. Multiple sequence alignments were made for each single copy ortholog using MAFFT v.7.487 default parameters (Katoh & Standley, 2013). Unrooted gene trees were produced via IQ-TREE v.1.6.12 (Nguyen, Schmidt, Von Haeseler, & Minh, 2015) using 1000 ultrafast bootstrap approximations (Minh, Nguyen, & von Haeseler, 2013). Substitution models for each gene tree were determined via IQ-TREE’s built-in ModelFinder (Kalyaanamoorthy, Minh, Wong, von Haeseler, & Jermiin, 2017). Using these gene trees, we estimated the species tree with ASTRAL-Pro—a multispecies coalescent (MSC) summary method that accounts for orthology and paralogy (Zhang, Scornavacca, Molloy, & Mirarab, 2020). We then used ASTRAL-III (Zhang, Rabiee, Sayyari, & Mirarab, 2018) to gather quartet frequencies and test for polytomies (Sayyari & Mirarab, 2018). The species tree plot was produced using R packages ape (Paradis & Schliep, 2018) and treeio (Wang et al., 2019). To test for consistency across analysis types, we concatenated the gene alignments into one supermatrix using SequenceMatrix v.1.9 (Vaidya, Lohman, & Meier, 2011) and analyzed these data by Maximum Likelihood (ML) using IQ-TREE v2.2.0 (Minh et al., 2020) with 1000 ultrafast bootstraps. Within the supermatrix, we partitioned genes and determined substitution models according to ModelFinder, allowing each partition to evolve under its own evolutionary model (Chernomor, Von Haeseler, & Minh, 2016) (Fig. S2). Both the MSC and ML trees were rooted with *Fusarium abutilonis,* post-analysis.

### Gene Trees

Gene trees for historically relevant genes were conducted to further investigate the putative progenitors (Ekanayake et al., 2013). Blast indices were made for genes tubB, tefA, and perA (supp. Fig. 4). The genomes of sexual, non-hybrid *E. baconii* (GCA_023650635.1)*, E. bromicola* (GCA_023658445.1)*, E. festucae* (GCA_003814445.1)*, E. typhina* (GCA_023658615.1)*, E. typhina* subsp*. clarkii* (GCA_021378265.1)*, and E. amarillans* (GCA_024072835.1) were downloaded from NCBI as the most likely putative progenitors genomes to *Epichloë* hybrid species. *TefA, tubB* and *perA* gene sequences predicted by Blast were extracted from strains 1033 and 212 as well as from the non-hybrid genomes using Samtools v. 1.16 (Camacho et al., 2009; Danecek et al., 2021). *Fusarium solanis-melongenae* (CP090031.1 and CP090032.1) was included to serve as an outgroup for *tefA* and *perA* gene trees. A *tubB* gene was not found for *F. solani-melongenae* on the Blast webserver, so *F. solani* (OP294987.1) was used as the outgroup for the *tubB* gene tree*. F. solani* and *F. solani-melongenae* are members of the same species complex (Xie et al., 2022). Multiple sequence alignments were made for each gene using MAFFT v. 7.487 using --adjustdirectionaccurately -- nuc --localpair –maxiterate options (Katoh & Standley, 2013). Unrooted gene trees were produced via IQ-TREE v.1.6.12 (Nguyen et al., 2015) using 1000 ultrafast bootstrap approximations (Minh et al., 2013). Substitution models for each gene tree were determined via IQ-TREE’s built-in ModelFinder (Kalyaanamoorthy et al., 2017). Using these gene trees, we created a consensus gene tree with ASTRAL-Pro v. 1.10.1.3 with default parameters. (Zhang et al., 2020). Gene trees were rooted and visualized in FigTree v.1.4.4 (http://tree.bio.ed.ac.uk/software/Figtree/). To test for consistency, we concatenated the gene alignments into one supermatrix using SequenceMatrix v.1.9 (Vaidya et al., 2011) and analyzed these data by Maximum Likelihood (ML) using IQ-TREE v2.2.0 (Minh et al., 2020) with 1000 ultrafast bootstraps. Within the supermatrix, we partitioned genes and determined substitution models according to ModelFinder, allowing each partition to evolve under its own evolutionary model (Chernomor et al., 2016) (Fig. S3). Both trees were rooted with the respective *Fusarium* species, post-analysis.

### AT/GC and Repeat Analysis

Gene prediction was carried out by BRAKER (Brůna, Hoff, Lomsadze, Stanke, & Borodovsky, 2020; Hoff, Lomsadze, Borodovsky, & Stanke, 2019). The BRAKER annotations were fed into OcculterCut to determine the AT-rich regions (Testa et al., 2016). A custom repeat database was built with RepeatModeler v. 2.0.2. from each genome (Smit & Hubley, 2008-2015). The MITE database from Fleetwood et al. (2011) was added to each custom database. RepeatMasker v.4.1.1 was then run with the updated custom database (Smit, Hubley, & Green, 2013-2015). BRAKER gene annotations and OcculterCut regions were visualized on the genome assemblies using Geneious Prime (“Geneious Prime 2023 “).

### Alkaloid Genotyping

Alkaloid gene indices were built manually by searching for alkaloid genes on Uniprot.org from *E. coenophiala* and extracting the sequences from Genbank.org. If no genes were present for *E. coenophiala,* a closely related *Epichloë* species were utilized. Sequences used to build indices can be found in table S5.3. Blast was then used to create the indexes and then to find significant hits for each index. Blastn megablast was used and a max of 5 alignments for each query-subject pair was set. Blast work was carried out with version BLAST+/2.12.0-gompi-2020b (Camacho et al., 2009). A gene was recorded as present when a significant hit was found.

### Dot Plot Alignments

All alignments were carried out using MUMmer v. 4.0 nucmer command with a word length of 30 (Marçais et al., 2018). The show-coords command was used on the delta file and visualized using the R script dotPlotly with varying filter specifications (RStudio, 2022). Self-alignments were made for each strain assembly to determine the presence of sub-genomes. Strains were aligned against each other to determine genomic collinearity. Genome fragmentation post-hybridization was determined by first aligning putative progenitor genomes *E. typhina* and *E. baconii* (Fig. S2A), and highly collinear chromosomes of *E. typhina* chromosome 4 and *E. baconii* chromosome 3 were identified. The chromosomes were then aligned back to each strain assembly. Additionally, each progenitor’s whole genome was aligned to both strain assemblies (Fig. S2B-E).

Further investigations into potential progenitor-strain alignments were done using minimap2 with the -asm option for aligning different species (Li, 2018). To manually inspect alignments, the output was loaded into IgV (Robinson et al., 2011). The BRAKER gene annotations were also loaded into IgV to compare genome alignments to gene-rich regions.

## RESULTS

### Genome assemblies

PacBio HiFi sequencing yielded high-quality reads. Quality assessments show that high-quality genomes assemblies were produced (table 5.1). The 1033 assembly has a genome size of ∼83.6 Mb, and the 212 assembly has a genome size of 87.6 Mb. The 1033 assembly has 11 telomere-to-telomere contigs and the 212 assembly has 9 telomere-to-telomere contigs, suggesting these contigs represent a full chromosome. The 1033 assembly overall has 14 contigs that contain start telomere motifs and 11 contigs that contain end telomere motifs. The 212 assembly overall has 12 contigs that contain start telomere motifs and 13 contigs that contain end telomere motifs. Centromere motifs are not identified for *Epichloë* species are therefore could not be utilized for analysis. Each genome assembly produced many contigs, however, only a fraction of the contigs produced represent the majority of the sequence length (Fig. 2). The BUSCO score shows > 85% duplication for both assembled genomes. Visual assessment of the self-alignment dot plots of the strains shows a single contiguous alignment, indicating no substantial sub-genomic collinearity is present (Fig. 3).

**Figure 2.**
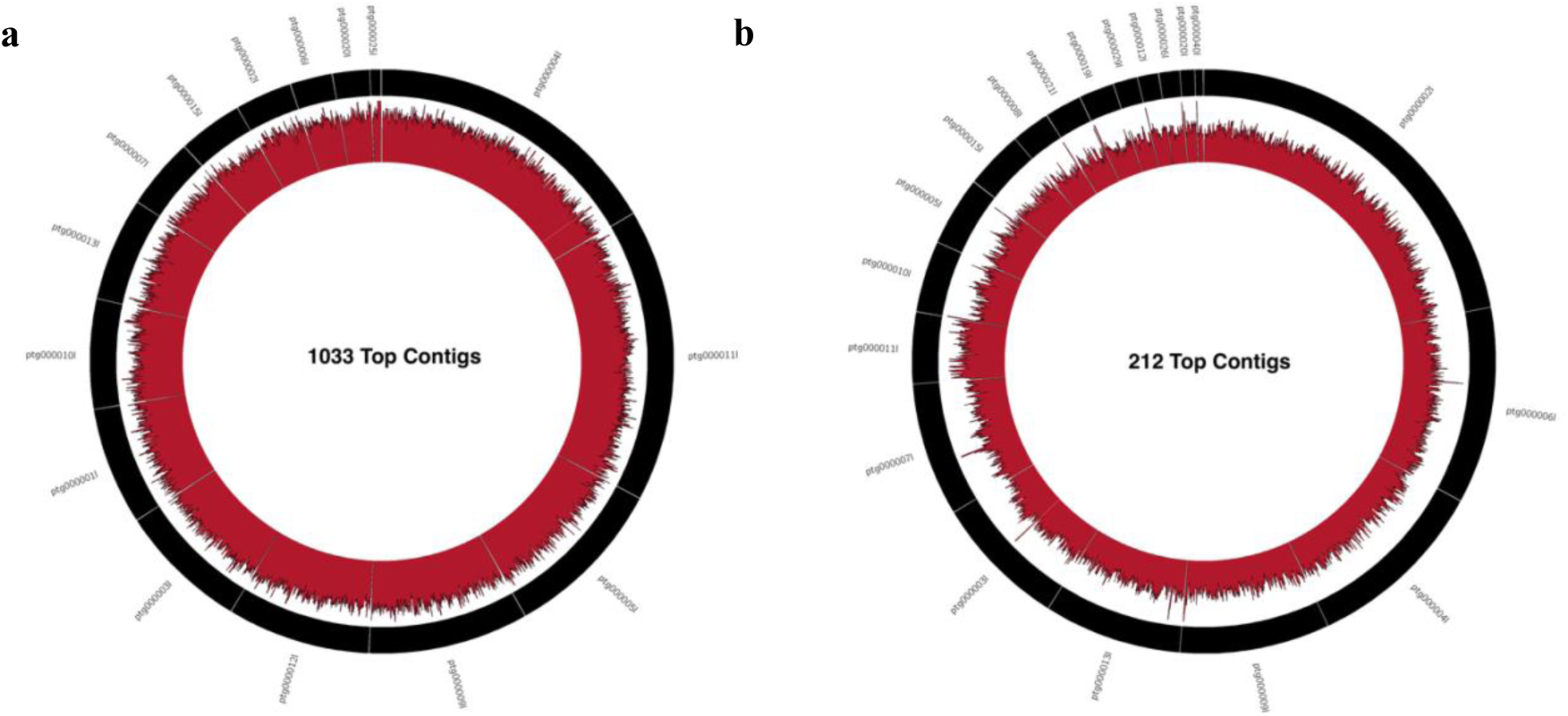
Base sequence coverage histogram for top contigs in the assemblies of a) strain 1033 and b) strain 212, where contigs shown represent a high percentage of total genome length (90.75% and 97.08%, respectively). Histogram maximum is 250 and minimum is 0.

**Figure 3.**
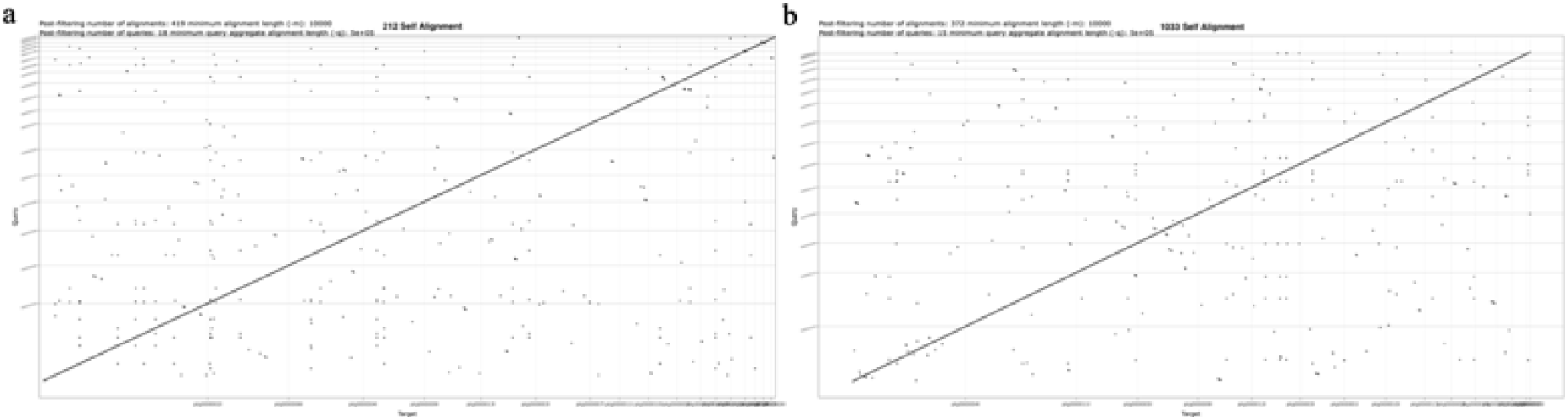
**Self-alignment dot plots for a) *Epichloë* strain 212 and b) strain 1033.**

### Progenitor Identification

Using Orthofinder we identified 109 single-copy genes shared across all ten samples included in phylogenomic analyses. Species relationships in the MSC summary tree had high local posterior probability (PP) support (PP=1) for all except the clade containing *E. bromicola, E. typhina,* and *Epichloë* strains 1033 and 212 (PP=0.52), and the smaller clade with *E. typhina* and *Epichloë* strains 1033 and 212 (PP=0.93) (Fig. 4). The branch leading to the clade containing *E. bromicola, E. typhina,* and *Epichloë* strains 1033 and 212 was relatively short and, according to the ASTRAL polytomy test, we cannot reject the null hypothesis that this branch should be replaced by a polytomy (*p-value*=0.59). This is also supported by the conflicting quartet frequencies at that node (q1=0.36, q2=0.35, q3=0.29) (Fig. 4). However, we can reject the null for all other branches in this tree (*p-value*=0). Regardless, the two *Epichloë* strains from this study are most closely related to *E. typhina* and *E. typhina* subsp. *clarkii,* which was also supported in the ML concatenated analysis (ultrafast bootstrap support=100). Tree topologies were mostly congruent between the ML and MSC analyses in that they both showed high support for the clade containing *E. festucae, E. baconii,* and *E. amarillans*, as well as the clade with *E. typhina, E. typhina* subsp. *clarkii* and *Epichloë* strains 1033 and 212, but could not confidently resolve the relationship between these two larger clades and *E. bromicola*.

**Figure 4.**
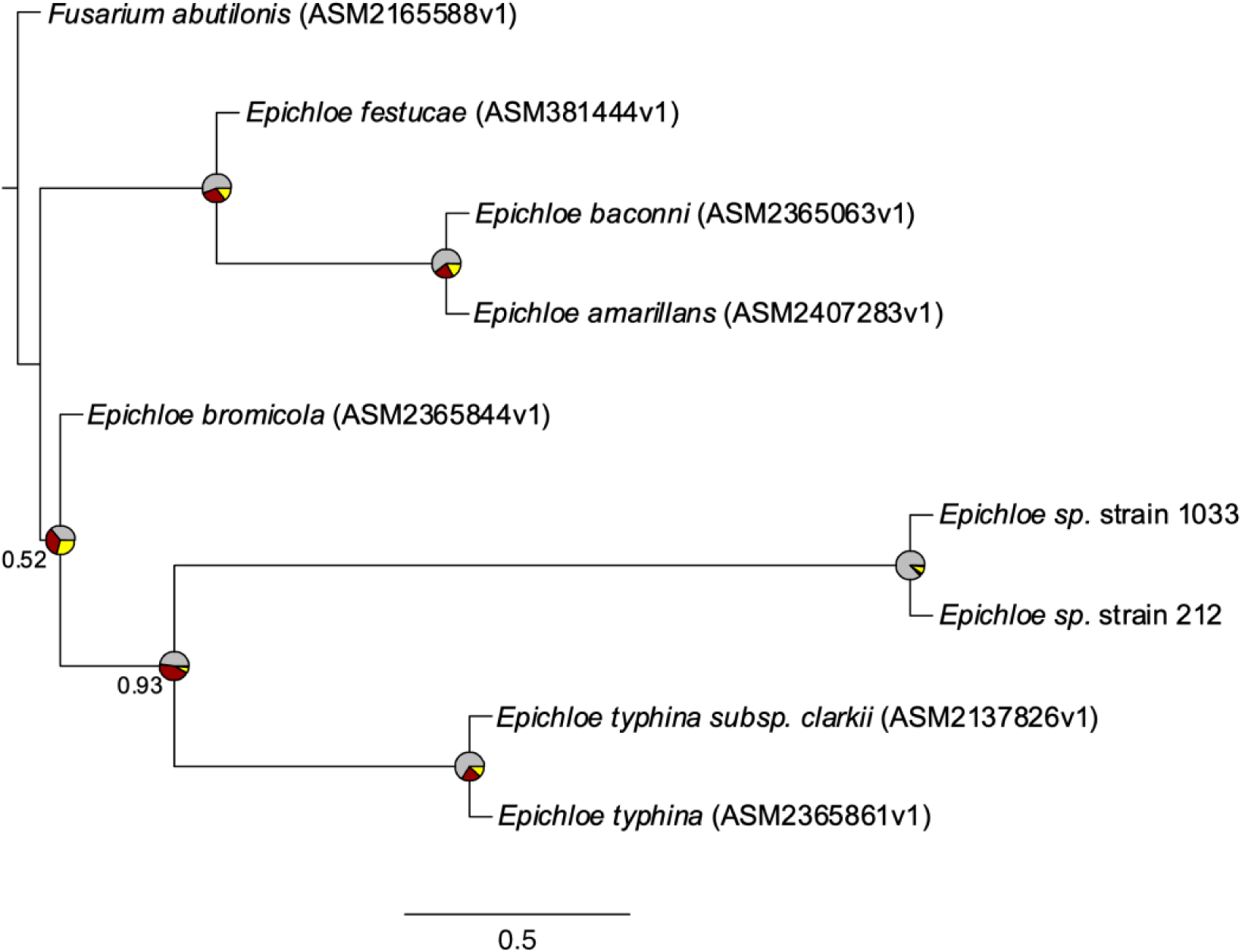
Rooted species tree showing relationships between the two *Epichloë* genomes assembled in this study (i.e., strain 1033 and 212) with those from previous studies. Pies are gene tree quartet frequencies supporting the main topology shown here (grey) as well as second (red) and third (yellow) alternatives. Node support values are local posterior probabilities and are only shown if < 1. Branch lengths and scale bar (bottom) represent coalescent units. Terminal branch lengths were not estimated but are illustrated here as pseudo branch lengths.

BLAST searches of gene homeologs predicted by BUSCO repeatedly suggest that *E. typhina* and *E. baconii* are the homoeologous gene donors of strain 212 (Supp. table 2 and 3). The gene tree made with nine randomly chosen genes, including both homologs found in the hybrid strains and the genes of their putative progenitors, shows perfect local posterior support for strain homeologs being diverged from their assigned progenitor species (Fig. 5). Based on these findings, we can conclude that *E. typhina* and *E. baconii* are the progenitor species of both strains.

**Figure 5.**
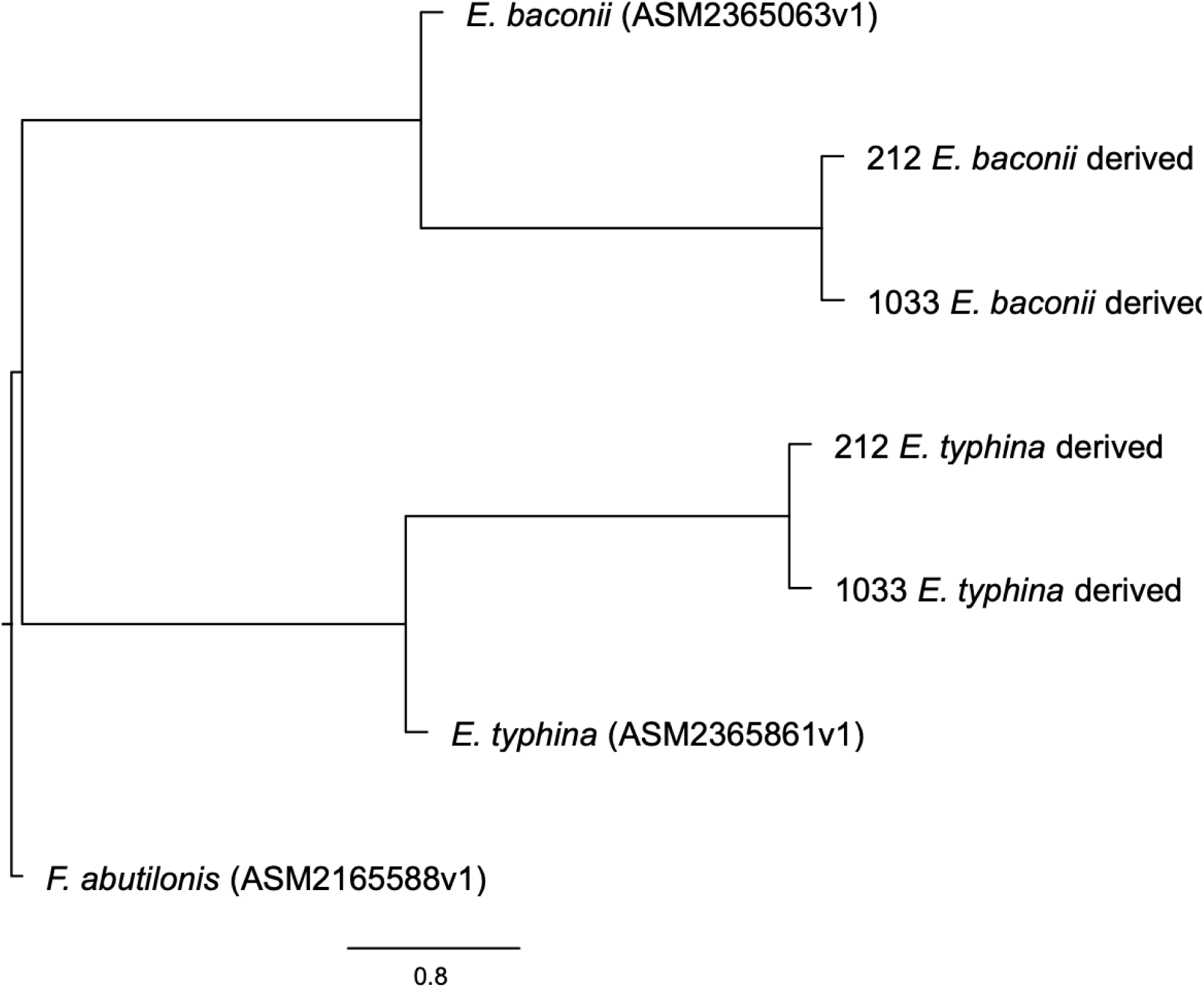
Rooted nine-gene tree that represents the putative progenitor genomes *E. typhina* and *E. baconii* and the gene homeologs from the strain assemblies (i.e., 212 and 1033) that were donated by each progenitor. Local posterior support was perfect for every node.

### Species characterization

We attempted to decipher the species of the fungi strains sequenced by analyzing putative chemotype, ploidy, progenitor species, and blast searches of canonical housekeeping and alkaloid genes. Both strains are diploid and are therefore likely one of the *E.* sp. FaTG species. Using a custom BLAST index, we were able to identify the presence or absence of all alkaloid biosynthesis genes in strains 1033 and 212. We found that the 212 assembly contains *lolF, lolP, lolT, lolN,* and *perA*. We can assign a tentative chemotype of loline-producing and peramine-producing to strain 212. The 1033 assembly contained alkaloid genes *lolF, lolP, lolT, lolN, perA,* and *lpsA.* 1033, therefore, likely produces lolines, peramine, and possibly ergot alkaloids. A similar method was employed to identify mating type genes in both strains. Both strains contained significant hits for *mtAA* and *mtAB* and only minor sequence hits for *mtAC*. Blast results indicate strong evidence of both strains being either *E.* sp. FaTG-3 or *E.* sp. FaTG-5, with multiple genes matching with 100% or very close to 100% identity and query coverage (table 2).

**Table 1.** Assembly statistics for strains 212 and 1033.

| Strain | Total Length | N50 | # of Contigs | BUSCO Completeness | BUSCO Duplication |
| --- | --- | --- | --- | --- | --- |
| 212 | 87,626,395 | 6130774 | 298 | 97.2% | 87.9% |
| 1033 | 83,594,297 | 6036110 | 344 | 97.4% | 86.7% |

**Table 2.** Predicted gene homeologs from each *Epichloë* strain’s top match when searched using NCBI BLAST.

| Strain | Gene Searched | Top BLAST result | Query coverage (%) | Identity (%) |
| --- | --- | --- | --- | --- |
| 1033 | *tubB* homeolog 1 | *Epichloë* sp. FaTG-5 | 100 | 100 |
| 1033 | *tubB* homeolog 2 | *Epichloë* sp. FaTG-3 | 100 | 99.92 |
| 1033 | *tefA* homeolog 1 | *Epichloë typhina* subsp. *typhina* | 99 | 99.5 |
| 1033 | *tefA* homeolog 2 | *Epichloë baconii* | 99 | 97.34 |
| 1033 | *perA* homeolog 1 | *Epichloë* sp. FaTG-3 | 100 | 100 |
| 1033 | *perA* homeolog 2 | *Epichloë* sp. FaTG-3 | 99 | 100 |
| 212 | *tubB* homeolog 1 | *Epichloë* sp. FaTG-5 | 100 | 99.92 |
| 212 | *tubB* homeolog 2 | *Epichloë* sp. FaTG-3 | 100 | 99.92 |
| 212 | *tefA* homeolog 1 | *Epichloë typhina* | 100 | 98.42 |
| 212 | *tefA* homeolog 2 | *Epichloë baconii* | 99 | 97.22 |
| 212 | *perA* homeolog 1 | *Epichloë* sp. FaTG-3 | 100 | 99.91 |
| 212 | *perA* homeolog 2 | *Epichloë typhina* | 100 | 99.27 |

### Alignments

Figure 6a shows the chromosome alignment of *E. typhina* chromosome 4 and *E. baconii* chromosome 3. These chromosomes are highly collinear. Figures 6b and 6c show how this collinearity is disrupted when *E. typhina* chromosome 4 and *E. baconii* chromosome 3 were aligned to strain assemblies 212 and 1033. While there are regions of alignment between the progenitor genomes and the strain assemblies, there is not the same level of collinearity as seen between the progenitor genomes. The strain to progenitor alignments utilized the top contigs from each assembly (Fig. 2). The 212 strain contigs that are shown in Fig. 6b range from ∼17.8Mb (ptg000002l) to ∼1.5Mb (ptg000019l) in size. The 1033 strain contigs that are shown in Fig. 6c range from ∼12.6 Mb (ptg000004l) to ∼3Mb (ptg000007l) in size. The assemblies of 212 and 1033 were also aligned and show significant fragmentation with small regions of high alignment but in a seemingly random manner (Fig. 6d).

**Figure 6.**
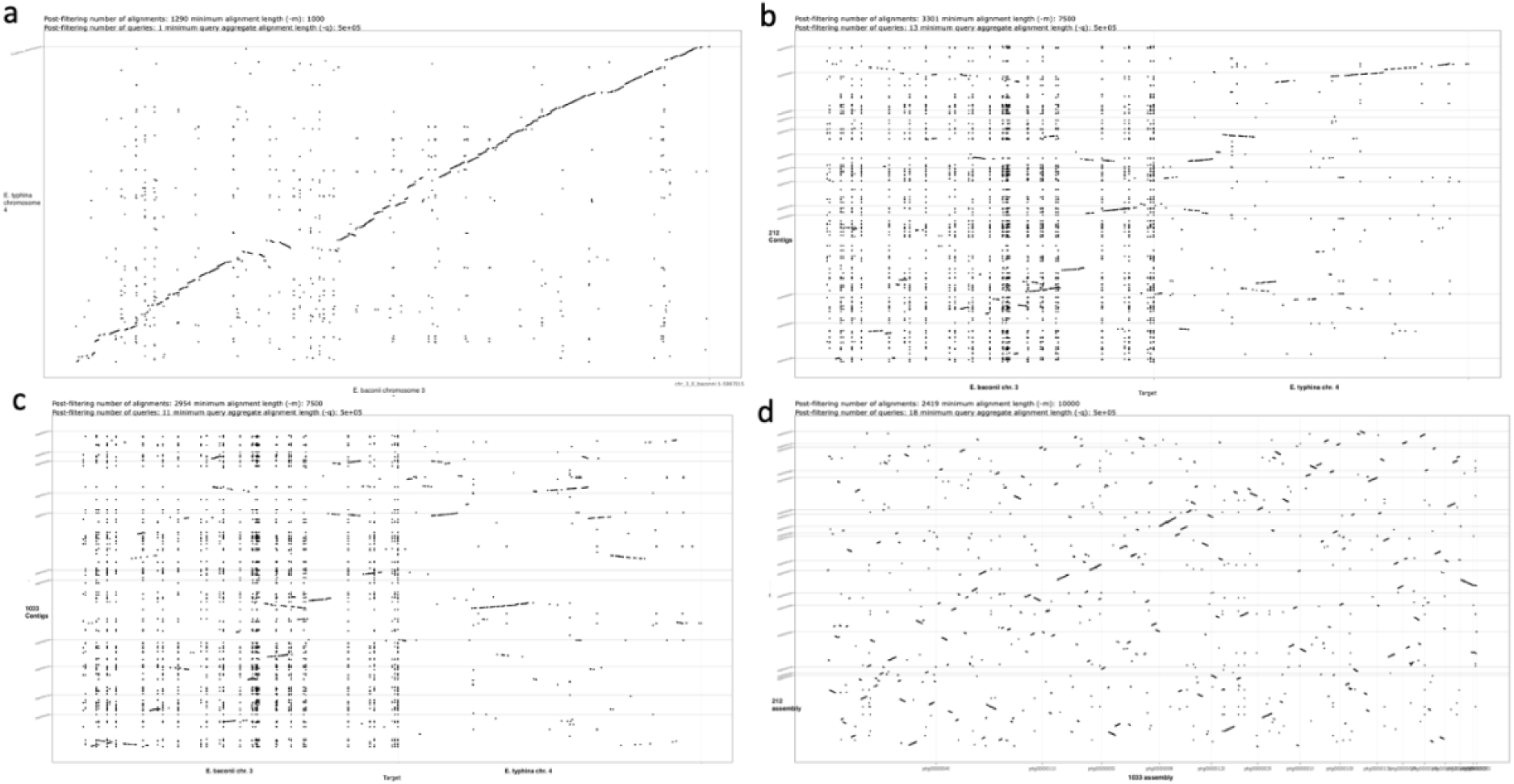
Alignment dot plots showing a) highly collinear alignment of *E. typhina* chromosome 4 to *E. baconii* chromosome 3, b) *Epichloë* strain 212 assembly to *E. typhina* chromosome 4 and *E. baconii* chromosome 3, c) *Epichloë* strain 1033 assembly to *E. typhina* chromosome 4 and *E. baconii* chromosome 3 and d) 1033 assembly aligned to the 212 assembly.

### AT/GC & Repetitive element content

Both assemblies were found to be highly composed of AT-rich regions (table 3). The strain assemblies were found to be consistent with past work on *Epichloë* with very clear blocks of AT-rich regions and GC-rich regions. Genes are not found in the AT-rich region of the genome.

**Table 3.** AT-rich region information for the genome assemblies of *Epichloë* strains 212 and 1033.

| Strain | AT Rich % | Mean Length of AT Rich Region (kbp) |
| --- | --- | --- |
| 212 | 43.9% | 34.2 |
| 1033 | 41.9% | 31.1 |

Repeat analysis showed a high percentage of each genome assembly being composed of repetitive elements. Strain 212’s genome assembly is 45.32% RE. Similarly, strain 1033’s genome is 43.01% RE (Fig. 8a). The repetitive elements of each assembly were mainly retrotransposons Gypsy/DIRS1 and Ty1/Copia (Fig. 7a). A high frequency of Miniature Inverted Transposable Elements (MITEs) was also found in each genome (Fig. 8B).

**Figure 7.**
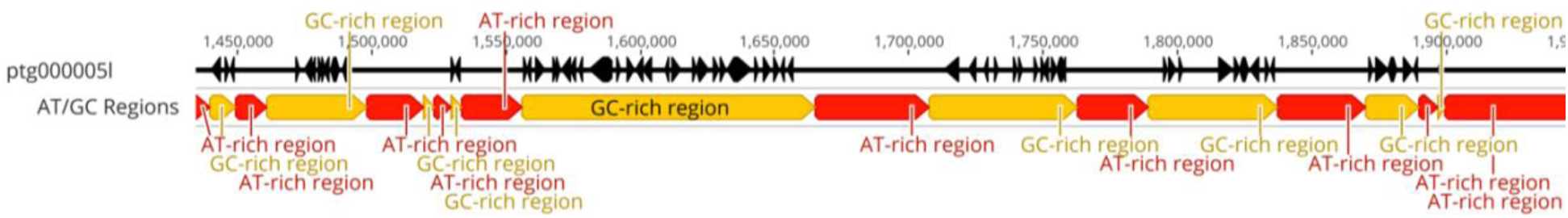
Example of AT-rich regions being absent of genes in *Epichloë* strain 1033. Genes are represented by black arrows. Red regions represent AT-rich regions. Yellow regions represent GC-rich regions.

**Figure 8.**
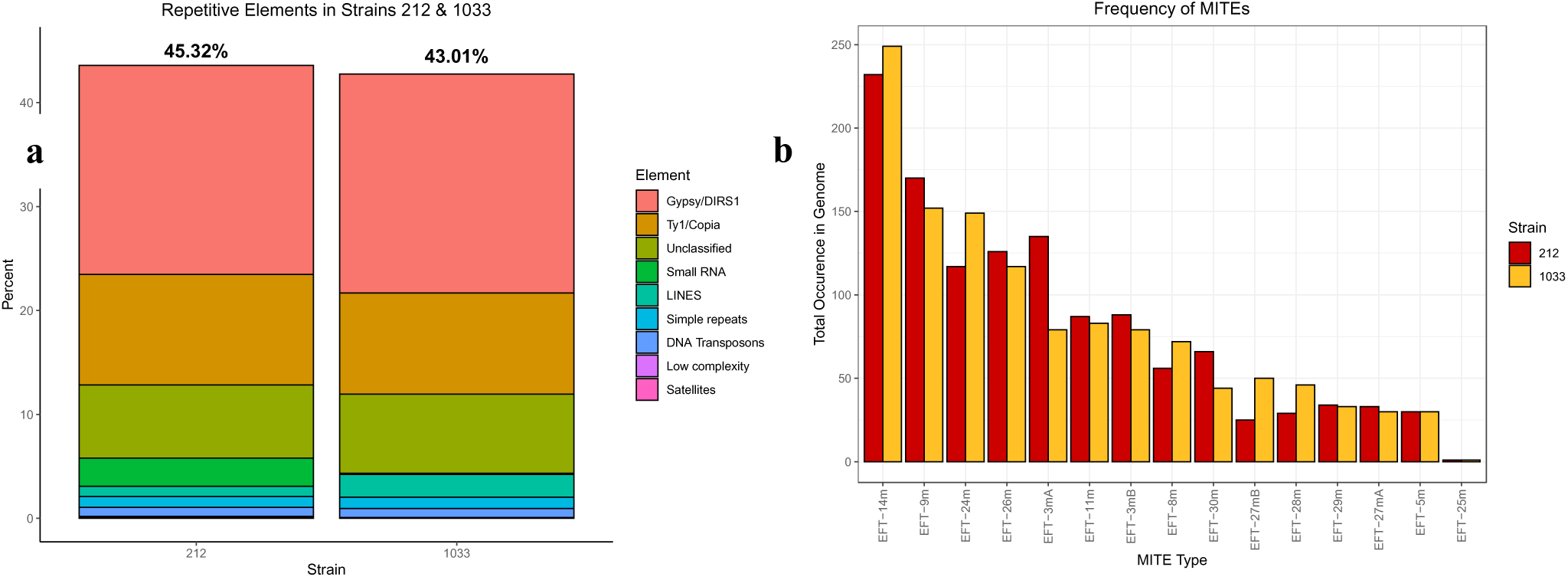
Repetitive element analysis of *Epichloë* strains 1033 and 212. a) Percent of repetitive element types in each assembled genome. b) The frequency of occurrence of different types of MITEs previously identified as abundant in *Epichloë* species in each assembled genome.

## DISCUSSION

### Discussion

In this study, we report the production of high-quality genome assemblies for two previously un-sequenced hybrid *Epichloë* strains derived from different tall fescue morphotypes. Hybrid *Epichloë* species have long lacked high-quality whole-genome sequences due to their complex genomic structures, making our assemblies a significant contribution to tall fescue endophyte research. The assemblies presented here contain several telomere-to-telomere contigs, suggesting the presence of full chromosome assemblies. However, full chromosome scaffolding would require Hi-C data. Despite this, the assemblies demonstrate the power of PacBio HiFi sequencing in generating highly contiguous sequences.

By aligning these assemblies with previously described species, we could tentatively assign the strains to species types based on existing species descriptions (Ekanayake et al., 2017; Takach & Young, 2014; Young et al., 2014). While *E. coenophiala* was the expected species type, the results suggest that only two progenitor species contributed to both strains. The projected chemotypes of strains 212 and 1033 align with *E. sp. FaTG-3* and *E. sp. FaTG-5*, respectively. *E. sp. FaTG-3* has *E. typhina* as a progenitor and a mating type of AA, and while discrepancies between our strain assemblies and previously described species exist, we confidently classify these strains as *E. sp. FaTG-3*-like.

We could identify only one probable progenitor in this study. Gene tree analyses, based on tefA and tubB, along with species/progenitor phylogenetic trees, suggest that *E. typhina* is a progenitor to both strains 212 and 1033. However, the identification of a second progenitor remains elusive. It is possible that the second progenitor is not present in the current sampling scheme or that the B gene copies from strains 212 and 1033 have evolved since hybridization (Fig. 5). Gene duplication cannot be ruled out, and the gene trees show that gene copies A and B from the strains are nearly identical, as evidenced by the lack of branch length (Fig. 5).

The genome sizes of strains 212 and 1033, 87.6 Mb and 83.5 Mb, respectively, align with the fusion of progenitor genomes. Non-hybrid *Epichloë* genomes typically range from 33.2 to 46.2 Mb (Quenu et al., 2022), with the *E. typhina*and *E. baconii* genomes used for comparison being 39.8 Mb and 39.2 Mb, respectively. The strains in this study show genome sizes that are approximately 8.6 Mb and 4.5 Mb larger than the combined progenitor genomes, suggesting the expansion of certain genomic regions. Repetitive elements likely contribute to this size discrepancy, as variation in non-hybrid *Epichloë* species is known to result from differences in transposable element (TE) content (Quenu et al., 2022). Non-hybrid *Epichloë* strains exhibit uncorrelated size variations, large chromosomal rearrangements, and variation in AT-rich regions, all while maintaining seven chromosomes. Similarly, the assemblies in this study have a comparable gene count but show significant rearrangements and fragmentation when aligned to each other. Both progenitor genomes contain approximately 7,500 genes (Quenu et al., 2022), and a fully duplicated hybrid genome would yield around 15,000 genes. The BUSCO analysis predicted 8,474 genes for strain 212 and 8,414 for strain 1033, which is substantially fewer than expected for full gene retention, suggesting significant gene loss or missing genes in the BUSCO Hypocreales database. However, gene predictions using BRAKER, which employs more robust parameters, yielded a higher count of around 10,600 genes for each strain (Table 3) (Hoff et al., 2019).

Dot plot alignments of the strain assemblies to their progenitor genomes and to each other provide strong evidence that the hybridization process resulted in extensive genome fragmentation and rearrangement, a hallmark of each hybridization event (Fig. 6). It is well documented that hybridization can lead to large-scale deletions, amplifications, and rearrangements of genomic regions until a stable genome is established (Steensels et al., 2021). These mechanisms likely occurred in strains 212 and 1033, resulting in the genomic upheaval observed. A striking feature of both assemblies is the high retention of gene copies, despite the extensive genomic changes. Moreover, each gene homolog remains highly similar to its corresponding progenitor gene (Supp. Table 2 & 3), suggesting strong selective pressure to retain both ancestral gene copies.

We also found a high frequency of repetitive elements, particularly MITEs, among the transposable elements (TEs) in both strain genomes. Previous studies align with this observation (Fleetwood et al., 2011), and it is well known that TEs can drive genome rearrangements. While MITEs and other TEs likely contribute to the observed fragmentation, the extent of the genomic upheaval may be driven by hybridization-specific mechanisms. It is possible that this genomic fragmentation is stochastic and may have a neutral effect on the fungi, as gene retention remains high. Alternatively, this upheaval could confer a selective advantage to the hybrid fungi, as the proximity of genes to TEs may influence gene expression (Winter et al., 2018), with certain genomic rearrangements potentially being beneficial.

Given the large-scale genomic disruption observed in hybrid *Epichloë*, reference-based genome assembly methods are likely unsuitable for these species. The extensive genomic turmoil observed here, even in highly similar strains, suggests that reference-based approaches would be ineffective. De novo assembly of hybrid *Epichloë* genomes, combined with graph-based methods for scaffolding and correction, would likely be more effective in addressing these challenges.

### Conclusion and Future Directions

Our findings provide strong evidence that parasexual interspecific hybridization causes profound genomic disruption in *Epichloë* strains, resulting in hybrid genomes that poorly align with either the progenitor or other hybrid genomes. Despite this, gene retention remains high, indicating that genomic changes occur without compromising the integrity of the genes. These results demonstrate that reference-based methods are not suitable for whole-genome analysis of hybrid *Epichloë* species. Expanding this study to include other hybrid *Epichloë* species would offer additional insight into the frequency and extent of genomic disruption following hybridization. Further studies investigating the transcriptomic and proteomic differences between hybrid *Epichloë* strains and their progenitors could provide valuable insights into the functional consequences of these genomic changes.

