## Supplemental Document for "Genomic Characterization of Novel Endophyte Strains from Tall Fescue Shows Genome Fragmentation Post-Hybridization"

Supplementary Table 1. Characterization of known *Epichloë* species and the samples characterized in this research. Efe = *E. festucae*, Ety = *E. typhina*, LAE = *Lolium-*associated endophyte, and Eba = *E. baconii.*

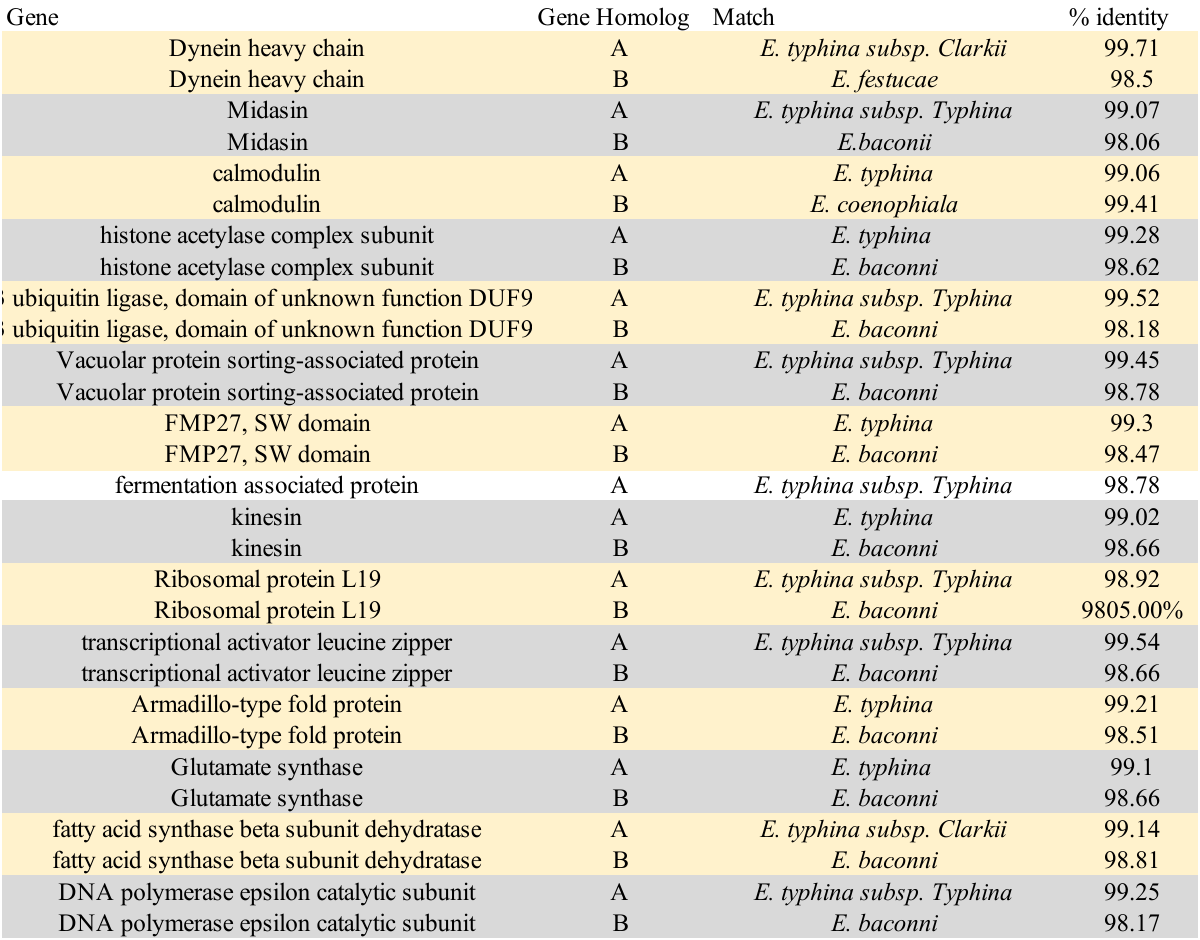

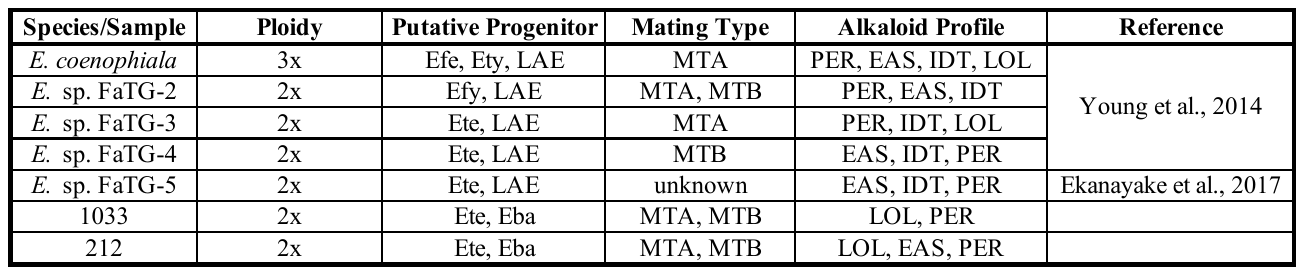

Supplementary Table 2. Strain 1033 gene blast top results for top scoring genes predicted by BUSCO.

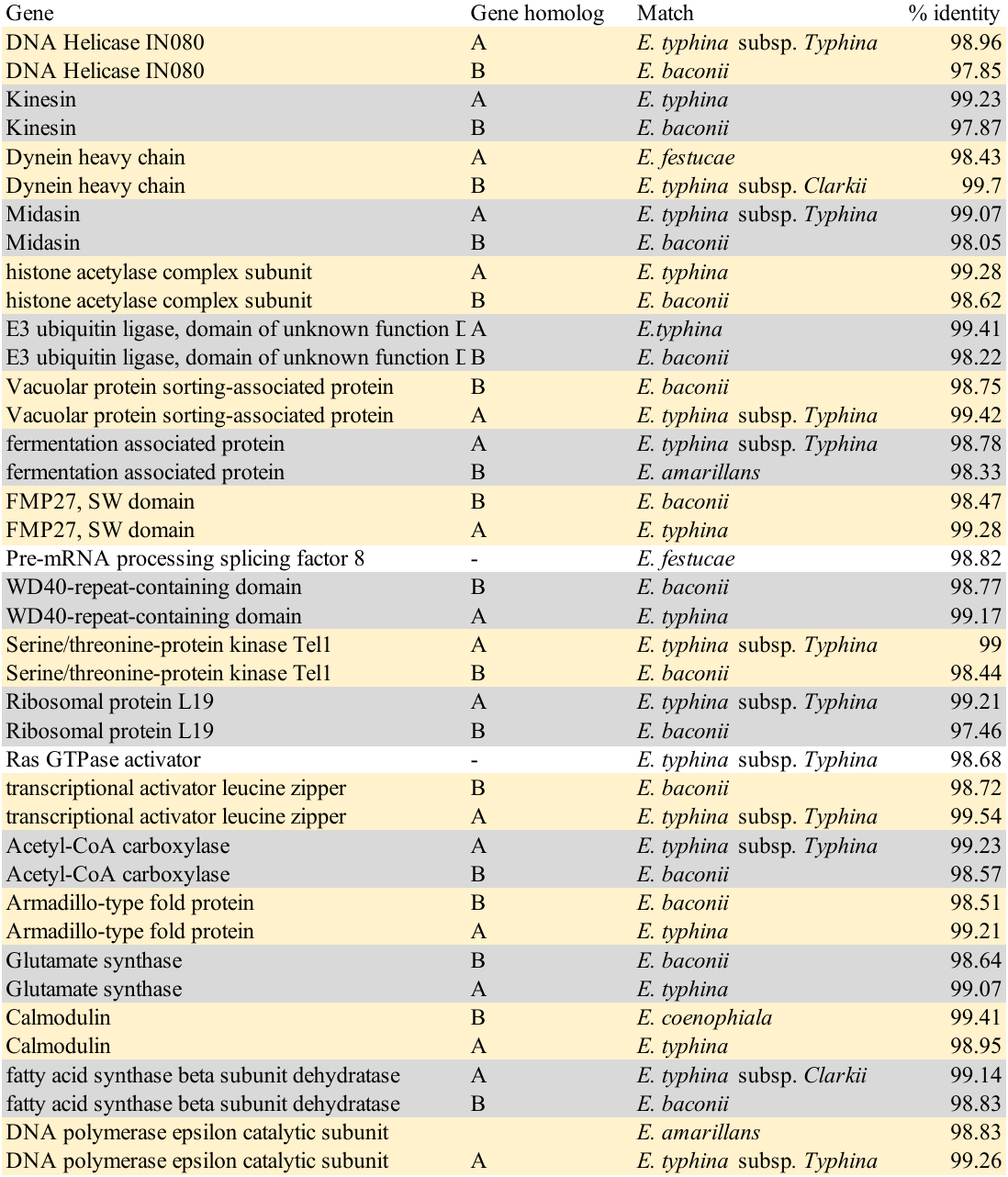

Supplementary Table 3. Strain 212 gene blast top results for top-scoring genes predicted by BUSCO.

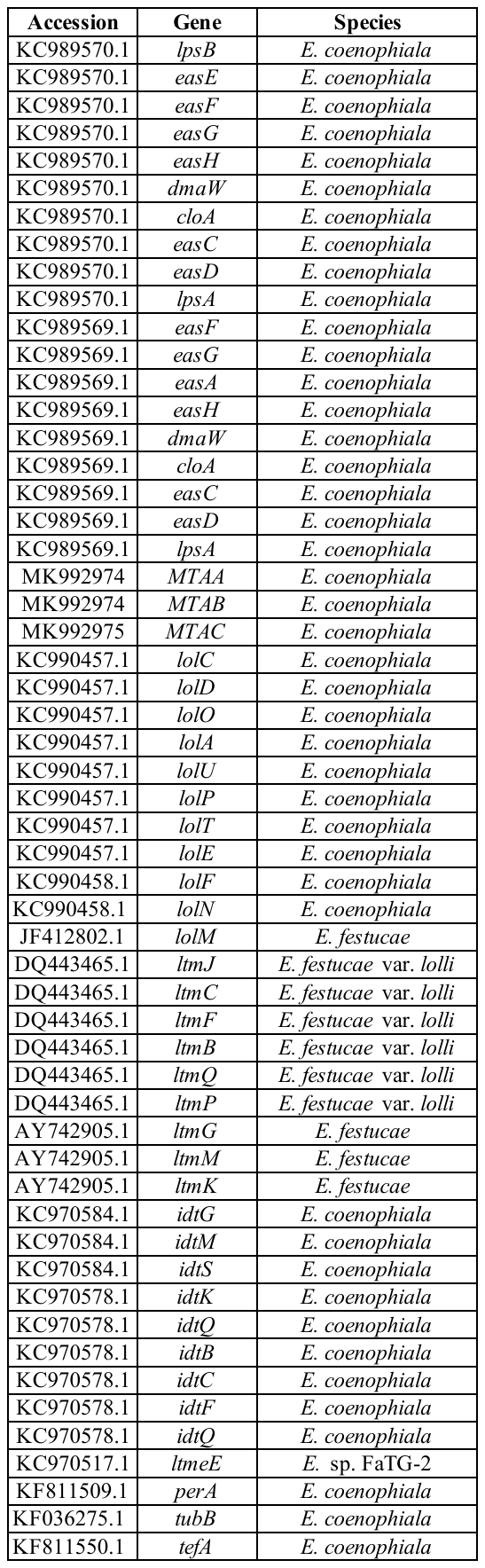

Supplementary Table 4. Genbank accessions used for BLAST indices.

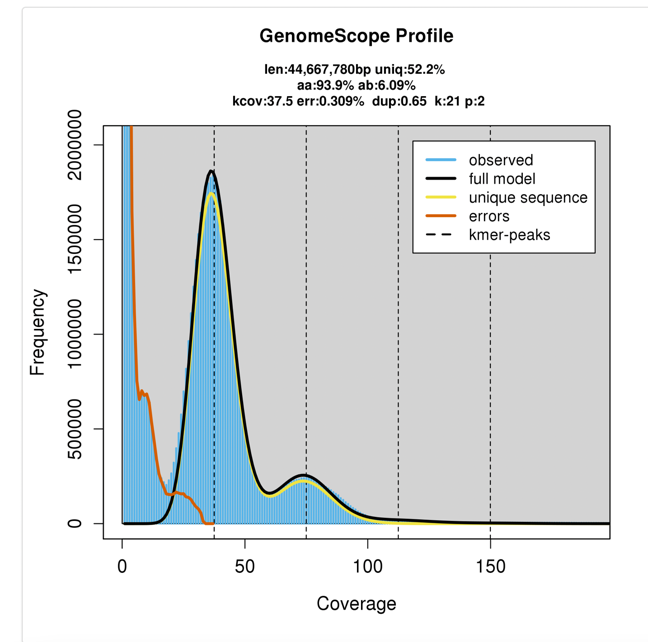

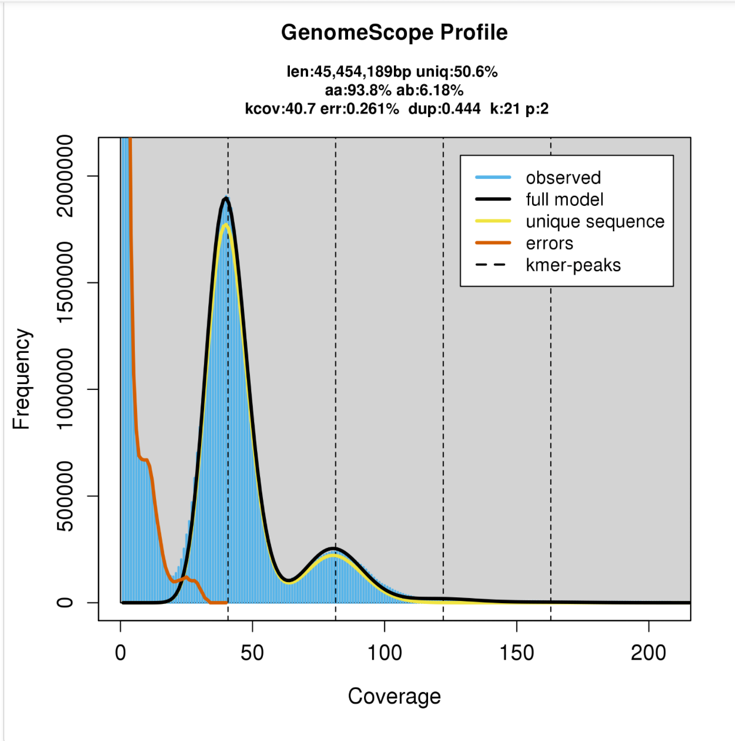

Supplementary Figure 1. K-mer histograms generated from GenomeScope 2.0 representing 21-mer count for the raw circular consensus sequences of strains A) 212 and B) 1033. K-mer histograms suggest that both strains are diploid and highly repetitive.

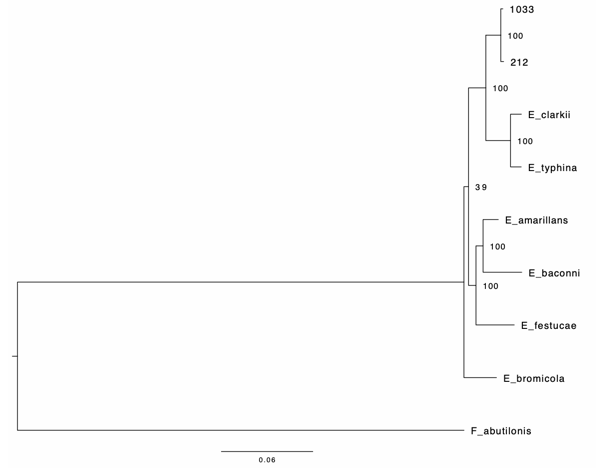

Supplementary Figure 2. Rooted maximum likelihood species tree of concatenated sequence data showing relationships between the two *Epichloe* genomes assembled in this study (i.e., strain 1033 and 212) with those from previous studies. Branch lengths and scale bar (bottom) represent coalescent units. Terminal branch lengths were not estimated but are illustrated here as pseudo branch lengths.

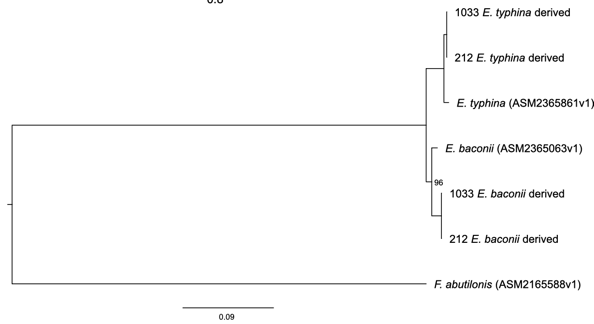

Supplementary Figure 3. Rooted maximum likelihood nine-gene tree of concatenated sequence data showing relationships between putative progenitor genomes *E. typhina* and *E. baconii* and the gene homeologs from the strain assemblies (i.e., 212 and 1033) that were donated by each progenitor. Local posterior support was perfect for every node except *E. baconii* clade where it is 96.

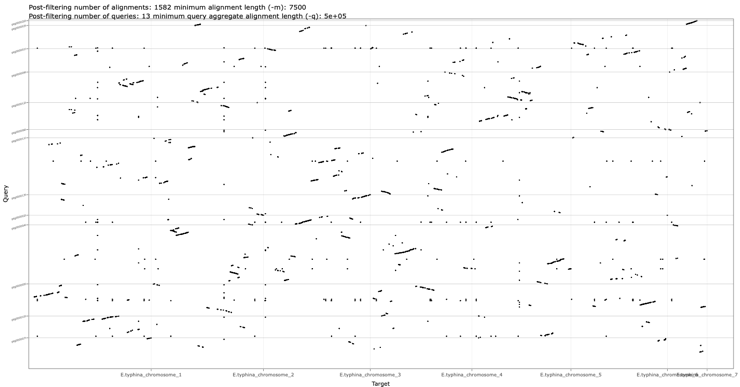

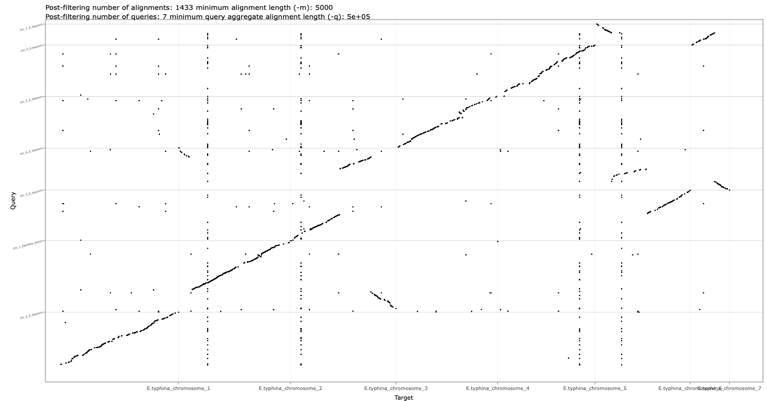

**B**

**A**

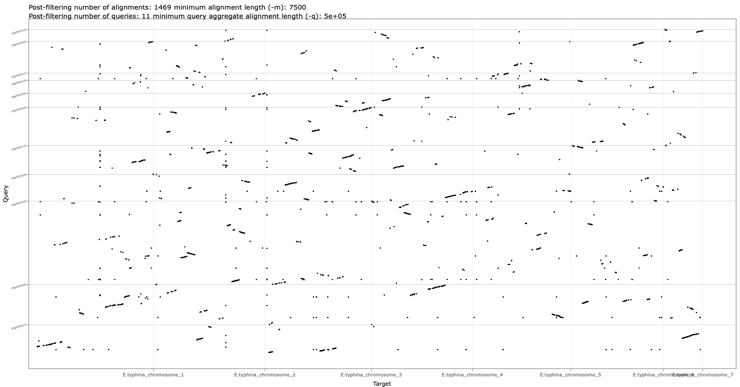

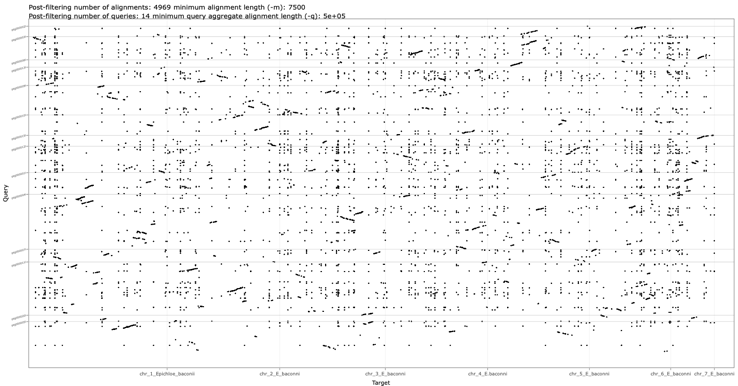

**C**

**D**

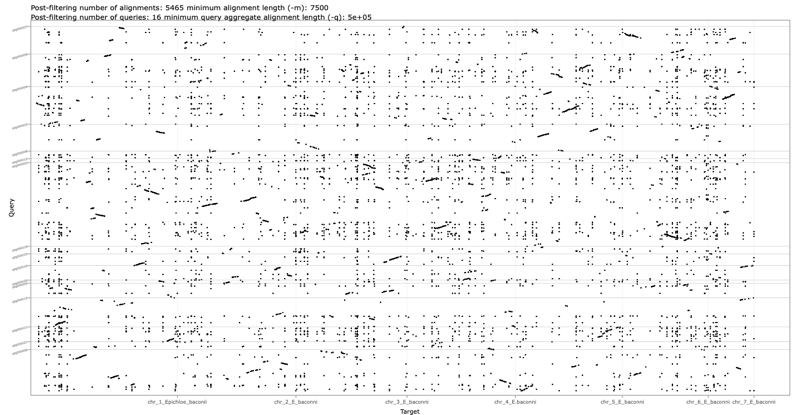

**E**

Supplementary Figure 4. Further alignment dot plots. A) full *E. typhina* and *E. baconii* genomes aligned. B) strain 1033 assembly aligned to the genome of *E. typhina.* C) Strain 1033 assembly aligned to the genome of *E. baconii*. D) Strain 212 assembly aligned to the genome of *E. typhina*. E) Strain 212 assembly aligned to the genome of *E. baconii.*
